# Too much, too young? Altered corticolimbic axonal innervation and resting state functional connectivity suggests sex-dependent outcomes in a rat model of early life adversity

**DOI:** 10.1101/700666

**Authors:** Jennifer A. Honeycutt, Camila Demaestri, Shayna Peterzell, Marisa M. Silveri, Xuezhu Cai, Praveen Kulkarni, Miles G. Cunningham, Craig F. Ferris, Heather C. Brenhouse

## Abstract

Adverse early experiences significantly alter behavioral and neural trajectories via aberrant brain maturation. Children with a history of early life stress (ELS) exhibit maladaptive behaviors and increased risk of mental illness later in life. Evidence in ELS-exposed humans identifies a role of atypical corticolimbic development; specifically, within amygdala-prefrontal cortex (PFC) circuits, and show precocially mature task-based corticolimbic functional connectivity (FC). However, the neurobiological substrates of such ELS-driven developmental changes remain unknown. Here, we identify putative neurobiological changes to determine the timeline of developmental perturbations following ELS in rats. Anterograde axonal tracing from basolateral amygdala (BLA) to pre- and infralimbic (PL, IL) PFC was quantified at postnatal days (PD)28, 38, and 48, along with anxiety-like behavior, in maternally separated (ELS) or control reared (CON) male and female rats. Resting state (rs)FC was assessed at PD28 and PD48 in a separate cohort. We report that ELS-exposed female rats show early maturation of BLA-PFC innervation at PD28, with ELS-related changes in males not appearing until PD38. ELS disrupted the maturation of rsFC from PD28 to PD48 in females, with enduring relationships between early rsFC and later anxiety-like behavior. Only transient ELS-related changes in rsFC were seen in male PL. Together, these data provide evidence that female rats may be more vulnerable to the effects of ELS via precocial BLA-PFC innervation, which may drive altered corticolimbic rsFC. These data also provide evidence that increased BLA-IL rsFC is associated with behavioral resiliency following ELS in female rats, providing mechanistic insight into the underlying etiology of adversity-induced vulnerability and resiliency.

## Introduction

Exposure to early life stress (ELS) leads to increased vulnerability to various psychiatric disorders across the lifespan (Callaghan and Tottenham, 2016; Hane and Fox, 2016; Krugers et al., 2017; Maccari et al., 2014; McEwan, 2008; Smyke et al., 2007). Males and females appear to be affected differently by ELS, with females more prone to developing disorders including anxiety and depression (Davis and Pfaff, 2014; Hammen et al., 2000; Heim et al., 2008). Importantly, these sequelae often emerge later in childhood or adolescence, providing an opportunistic window for intervention before psychopathology takes hold. Thus, development of effective intervention strategies requires biological and developmental targets specific to individuals based on factors including sex and timing of stress exposure (Lupien et al., 2009). Growing evidence suggests that ELS in humans leads to life-long changes in connectivity and/or functionality of limbic and cortical regions (Choi et al., 2009; Van Tieghem and Tottenham, 2018), with consequential deficits in emotion regulation and cognition (Tyrka et al., 2013). Human and animal studies highlight the importance of corticolimbic circuitry in affective behavior regulation, its disruption in mental disorders (Bangasser and Valentino, 2014; Herringa et al., 2013; Killgore et al., 2014), and alterations directly related to ELS (Kaiser et al., 2018). Specifically, Tottenham and colleagues illustrated that normative developmental changes in task-based functional connectivity (FC) between amygdala and medial prefrontal cortex (mPFC) are accelerated after ELS with associated changes in anxiety (Gee et al., 2013a) and are particularly evident in females (Dickie and Armony, 2008). Indeed, children institutionalized during the first two years of life in orphanages display precocial development of amygdala-mPFC FC, though the anatomical substrates for this accelerated connectivity remain unknown.

In rats, the basolateral amygdala (BLA) sends inputs to the mPFC and modulates anxiety-related behaviors (Felix-Ortiz et al., 2016), recall of emotionally salient information (St. McGaugh, 2004), decision-making (Onge et al., 2012), and goal-directed behavior (Schoenbaum et al., 2000). Notably, two subpopulations of BLA neurons project to different regions of the mPFC (Senn et al., 2014): projections to the dorsal (prelimbic; PL) region of the mPFC are active during threat-associated fear learning and expression, whereas projections from the BLA to the ventral (infralimbic; IL) region of the mPFC are active upon extinction of fear, or learning about safety signals. In typically developing male rats, BLA innervation of mPFC increases through adolescence (Cunningham et al., 2002), likely contributing to healthy maturation of threat and safety appraisal. Interestingly, BLA-mPFC axonal innervation has not yet been evaluated in females, nor has any study examined the effects of ELS on BLA-derived innervation of PL or IL.

Little is known about the interaction of sex and ELS on corticolimbic development (Herringa et al., 2013), despite striking sex differences in clinical time-course and symptomology of ELS effects (Martin et al., 2014; Wainwright and Surtees, 2002). Notably, typically developing females display earlier maturation of the PFC (Lenroot et al., 2007; Lenroot and Giedd, 2010). Identifying how ELS affects these sex-dependent trajectories is crucial to understanding sex differences in vulnerability and to develop individually targeted intervention strategies. Therefore, we used anterograde tracing to examine ELS effects on BLA-PFC innervation over development in male and female rats. We hypothesized that if heightened anxiety-like behaviors following ELS are associated with increased BLA-derived PFC innervation, then rats exposed to ELS via repeated maternal and peer separation will display anxiety-like behavior and increased BLA-PFC innervation that will be more robust in females.

While task-based FC illustrates coordinated responsivity to anxiety-provoking stimuli (Gee et al., 2013a), resting-state FC (rsFC) is excellent for probing the functional integrity of the amygdala-PFC circuit independent of task demands (Alarcón et al., 2015; Gabard-Durnam et al., 2016; Thomason et al., 2011a; 2011b). ELS effects on corticolimbic rsFC in humans are inconsistent, likely because of reliance on autobiographical questionnaires, different ages of measurement, and different ELS criteria. In rats, maternal separation results in early emergence of both adult-like fear learning based in fronto-amygdala circuitry (Callaghan and Richardson, 2011) and early amygdala structural maturation (Ono et al., 2008). Early maturation of corticolimbic connectivity likely has deleterious consequences because a sufficient degree of PFC immaturity during juvenility is critical for learning anxiolytic cues (safety signals) in adulthood (Yang et al., 2012). Therefore, we provide a back-translation to examine whether ELS-exposed rats display accelerated maturation of amygdala-PFC connectivity to parallel humans with a history of adversity, with increased BLA-PFC innervation as an anatomical substrate driving sex-specific developmental effects.

## Methods and Materials

### Subjects

For Studies 1 (BLA-mPFC innervation) and 2 (rsFC), timed-pregnant Sprague-Dawley rats (Charles River, Wilmington, MA) arrived at gestational day 15. Rats were housed under standard laboratory conditions in a 12h light/dark cycle (lights on at 0700h) temperature- and humidity-controlled vivarium with access to food and water ad libitum. Following birth (postnatal day [PD]0), litters were randomly assigned as either: control (CON), and left undisturbed with the exception of cage-changing twice/week and weighing (PD9, 11, 15, 20); or ELS via maternal and peer separation, as described previously (Coley et al., 2019; Farrell et al., 2016; Ganguly et al., 2019; Grassi-Oliveira et al., 2016; Wieck et al., 2013) and below. On PD1 litters were culled to 10 (+/-2) pups, maintaining equal ratio of male and female whenever possible, with one rat per litter assigned to each experimental group (i.e. age and sex) to avoid litter effects.

ELS pups were separated from dams and littermates in individual cups with home cage pine shavings in a circulating water bath (37oC) from PD2-PD10. At PD11-20, when body temperature is self-regulated, pups were individually separated into cages. Pups were separated for 4 hours each day (0900h-1300h) during which time pups were deprived of maternal and litter tactile stimulation and nursing, but not from maternal odor. ELS dams remained in their home cages but were deprived of their entire litters during separations. Pups were weaned at PD21 into same-sex mixed-litter pairs and left undisturbed until surgery/behavioral assessment for Study 1 (either PD21/28, 31/38, or 41/48; Fig1A). Separate cohorts were used for Study 2 – with treatment identical to Study 1 – and left undisturbed until behavioral assessment and rsFC (PD28, PD48), with subjects imaged at both ages. Experiments were performed in accordance with the 1996 Guide for the Care and Use of Laboratory Animals (NIH) and with approval from Northeastern University’s Institutional Animal Care and Use Committee.

**Figure 1.**
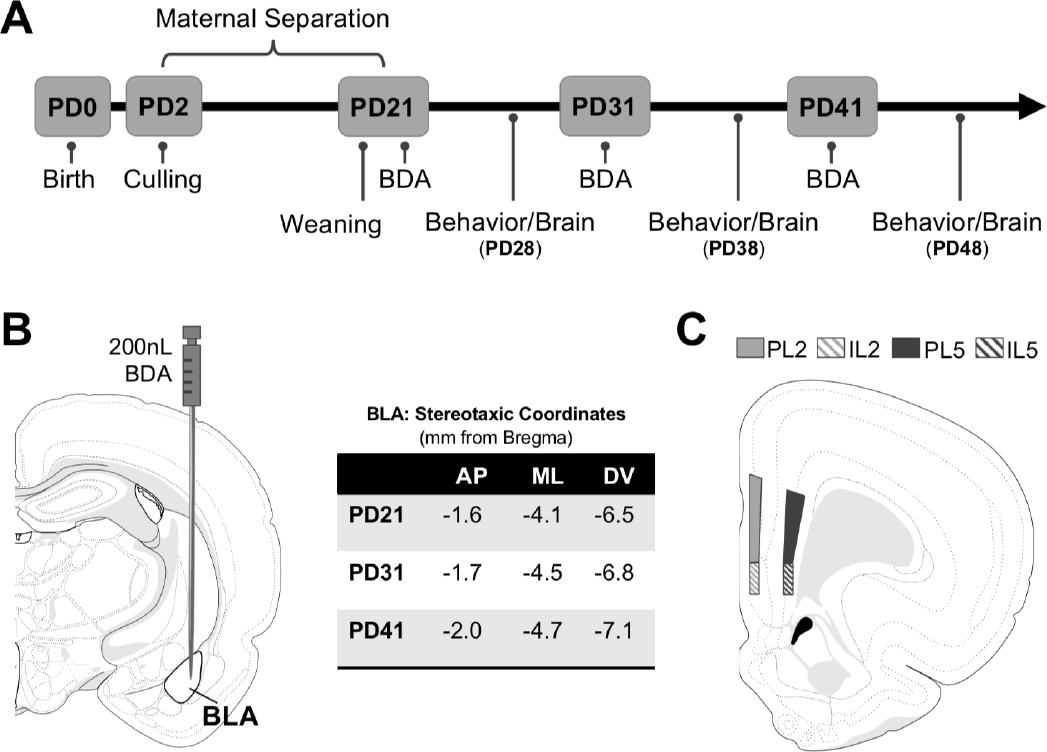
Timeline and methodology for study 1. (**A**) Methodological timeline for Study 1 indicating maternal separation (for ELS groups), weaning timeline, and surgical/behavioral milestones. Biotinylated Dextran Amine (BDA) microinfusions were performed at PD21, PD31, or PD41. Behavior (elevated plus maze; EPM) was performed at PD28, PD38, or PD48, and was followed by brain collection. (**B**) Stereotaxic coordinates for surgeries at each developmental time point and anatomical map of basolateral amygdala (BLA) injection site where 200nL of BDA was infused via Hamilton Neuros syringe. (**C**) Neuroanatomical map of quantified regions of BLA-PFC axonal innervation. Quantification was conducted via unbiased stereology within the prelimbic (PL) and infralimbic (IL) in layers 2 and 5. Atlas modified from Swanson (2018). AP (anterior-posterior); ML (medial-lateral); DV (dorsal-ventral)

### Study 1: BLA-PFC Innervation

#### Surgeries

Male and female rats from CON and ELS litters at PD21, PD31, or PD41 underwent stereotaxic injections of biotinylated dextran amine (BDA; Life Technologies NeuroTrace 10,000MW Anterograde Tracer Kit; reconstituted to 10% with phosphate buffered saline (PBS)) into BLA (Fig1B). For maximal uptake without excess bolus, 200nL (0.2μl) of BDA was injected into the BLA using a mounted 32-gauge Neuros syringe (Hamilton Company) attached to a microinfusion pump (11 Elite Nanomite; Harvard Apparatus).

Rats were first anesthetized with Isoflurane in an induction chamber before beginning surgical procedures. Surgical site was shaved, and the animal was secured via ear bars with top incisors positioned over a bite bar within a nose cone to provide continuous Isoflurane anesthetic during surgery. A subcutaneous injection of buprenorphine (0.03mg/kg body weight) was administered as a postoperative analgesic. Once sufficiently anesthetized, measured via lack of pedal reflex, a skin incision was made along the midline of the skull to visualize Bregma and Lambda. Dorsal-ventral (DV) coordinates for Bregma and Lambda were taken to ensure that the skull was level, and Bregma was used as a landmark to navigate to the position above the BLA, where a small hole was drilled into the right hemisphere of the skull, and the needle was slowly lowered into the BLA (Fig1B). DV depth was calculated from dura surface to account for individual differences in skull thickness. Once needle was lowered to the target, BDA was slowly infused at a rate of 40nL/min over the course of 5 minutes and left in place for an additional 5 minutes to allow for diffusion of BDA solution before being slowly retracted. The incision was sutured, and rats were returned to individual cages and allowed 6 days for recovery before behavioral testing, with all brain tissue collected one-week post-surgery.

#### Behavioral Assessment: Elevated Plus Maze

Elevated Plus Maze (EPM) performance was evaluated 6-7 days post-surgery in all animals. The apparatus was constructed of opaque Plexiglas with four radiating arms (50cm × 10cm) around a center square (10cm × 10cm), with 40cm-high walls surrounding two opposing arms, leaving the other two arms open. Between each animal 50% ethanol solution was used to clean the apparatus. Rats were acclimated to the testing room for 10min before testing began and were then placed in the center square of the apparatus under red light facing a closed arm. Behavior was recorded for 5 minutes by an observer blind to condition. Behavioral scoring included: four-paw entries into/time spent (seconds) in open and closed arms, number of arm crossings, and head dips.

#### Tissue Collection and Inclusion Criteria

7 days post-surgery, rats were deeply anesthetized with CO2 and transcardially perfused with ice-cold (0.9%) saline, followed by ice-cold 4% paraformaldehyde solution. Brains were extracted and stored in 4% paraformaldehyde solution for 1 week before being transferred to a 30% sucrose solution for 4 days for cryoprotection. All brains were sliced into 40μm serial sections on a freezing microtome (Leica), with serial sections placed in well plates filled with freezing solution for −20C storage. Sections taken for analysis included: PFC (+5.2mm through +2.5mm from Bregma) and BLA (−1.4mm through −3.6mm from Bregma) as outlined in Paxinos and Watson (1997). Serial sections were collected such that coronal sections within a single well were separated by approximately 240μm.

BLA-derived axonal innervation of the PFC, and BLA injection bolus for each animal, was visualized using diaminobenzidine (DAB) as indicated by the manufacturer (NeuroTrace BDA Kit, Life Technologies). One well for each animal was used for each brain region (PFC, BLA). Free-floating sections were washed 2×10min in PBS (7.4 pH) on an agitator before being transferred into Avidin solution (1:4000 in 0.3% PBS-T; Life Technologies NeuroTrace BDA Kit) overnight at 4oC on an agitator. Sections were then washed 3×5min in PBS and transferred into 0.05% DAB solution (Life Technologies NeuroTrace BDA kit) for 10 minutes for processing. Stained sections were washed in PBS, mounted on glass slides and coverslipped with DPX. Adjacent sections were randomly sampled from each experimental group (n’s=4/group) and stained with Cresyl Violet to determine average volume of the BLA based on age, sex, and condition. There were no significant differences based on sex or condition, thus volumes were collapsed across these to determine the average BLA volume per age (Supplemental Fig1).

#### Bolus Verification in the BLA and Inclusion Criteria

To determine the total volume of the BDA bolus within the BLA at the injection site, six DAB-stained sections containing BLA/BDA bolus were analyzed per animal (inter-slice interval of 240um). Volumetric analysis was conducted in StereoInvestigator (MBF Bioscience, Wilmington, VT) using a Cavalieri probe to estimate total bolus volume. Different markers were used to identify regions of the bolus that fell within versus outside of the BLA. Approximate percentage of BLA filled with BDA tracer was calculated by dividing the volume of bolus within the BLA by the average total BLA volume, specific to age, for each animal. Only subjects with a bolus filling greater than 60% of the average BLA volume (with no more than 15% of total bolus volume outside of the BLA) were included in analyses. See Fig2B,E for details on bolus size, location, and representative cases.

**Figure 2.**
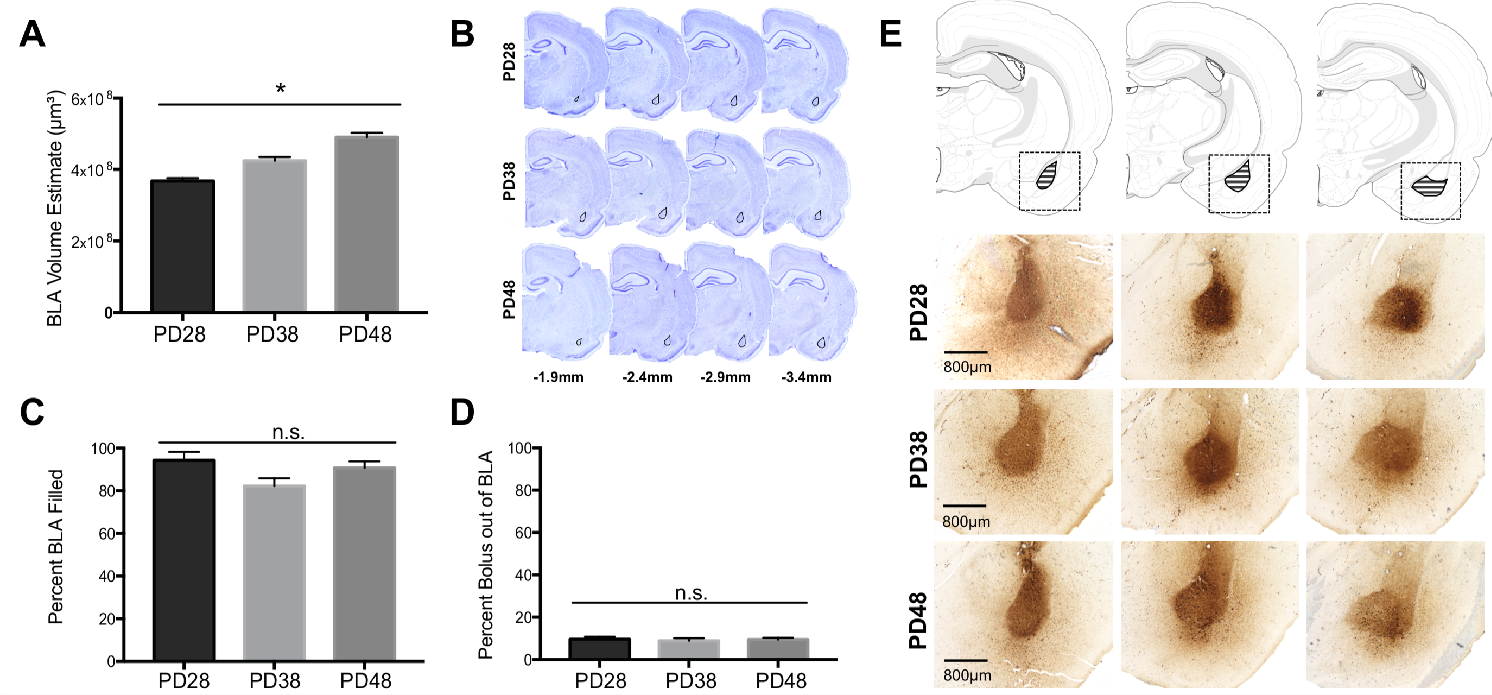
BLA volume and BDA anterograde tracer bolus sites. The average volume of the BLA significantly increased with age (**A**) as quantified via Cresyl Violet staining of adjacent tissue sections (**B**). As there were no significant differences as a function of sex or rearing condition on BLA volume estimates at each age group, data were collapsed for each age with a final number of n=16 per age group. Anatomical coordinates located below Nissl-stained sections (**B**) indicate the approximate distance (in mm) from Bregma. There were no significant differences in the percent of BLA filled by the BDA anterograde tracer bolus between ages (**C**), nor was there any significant difference in the amount of bolus located outside of the BLA structure (**D**). n = 6-9 per group (**C**; **D**) before collapsing within each age due to a lack of differences as a function of sex or rearing condition at each age (therefore, the number of data points per age group for analyses in **C** and **D** were 24-36). Panel (**E**) shows representative photomicrographs of BDA bolus across the extent of the BLA for each age examined. Atlas modified from Swanson (2018). \**p*<0.05; n.s. (non-significant)

#### Quantification of PFC Axonal Innervation

Three sections of each brain containing PFC (PL and IL regions: +4.0mm through +2.8mm from Bregma; 240um inter-slice interval) were used for StereoInvestigator analysis. BLA-derived axon fiber length was estimated via hemispheric SpaceBalls Probe in 4 regions per section: PL layer 2 (PL2) and 5 (PL5), and IL layer 2 (IL2) and 5 (IL5); Fig1C. Axonal fibers in focus over the probe (radius = 10μm) through the Z plane were identified at 60x oil immersion to determine estimated fiber length (Gundersen and Jensen, 1987; Gundersen et al., 1999; Mouton et al., 2002).

#### Statistical Analyses of Neuroantatomical and Behavioral Data

To determine effects of sex, rearing, and age on estimated total length of BLA-derived axonal innervation of the PFC, three-way ANOVAs were conducted within each region (PL, IL), and each layer (PL2, PL5, IL2, IL5). Layer-specific ANOVAs were analyzed with a Bonferroni alpha adjustment to correct for multiple comparisons, followed by post hoc individual comparisons with corrected p-values reported. Two-way ANOVAs were conducted separately for male and female groups to determine sex-specific effects. EPM performance was also analyzed with 3-way ANOVA and subsequent 2-way ANOVA within each sex. Significant main effects and/or interactions were followed up where appropriate with Bonferroni post-hoc analyses corrected for multiple comparisons. Trending effects were followed up with post-hoc analyses only with moderate-to-strong effect sizes determined via partial eta squared.

Separate linear regression analyses were conducted in male and female rats to determine whether there were relationships between behavior and BLA-PFC innervation. Fisher’s r-to-z transformations were used to assess any significant impact of sex or rearing on the strength of the relationships. Behavioral data collected from all subjects (regardless of whether they later met neuroanatomical inclusion criteria) was included in statistical analyses of EPM. However, all correlational analyses including behavioral data were performed only with the data from cases meeting neuroanatomical inclusion criteria. All statistical analyses were performed in either SPSS v25 (IBM) or PRISM 8 (Graphpad).

### Study 2: Resting State Functional Connectivity (rsFC)

MRI scanning was performed on male and female CON and ELS subjects at PD28 and PD48 in a Bruker BioSpec 7.0T/20cm USR horizontal magnet (Bruker, Billerica, MA) with a 20-G/cm magnetic field gradient insert (ID=12cm) capable of 120μs rise time. Rats were anesthetized and maintained at 1-2% isoflurane and oxygen with a flow rate of 1 L/min throughout scanning, with breathing rate (40-50 breaths per minute) carefully monitored by an investigator and anesthetic levels adjusted accordingly. Rats were scanned at 300 MHz using a quadrature transmit/receive volume coil built into the rat head holder and restraint system (Animal Imaging Research, Holden, MA). rsFC was acquired by gradient-echo triple-shot echo-planar imaging (EPI) pulse sequence with the following parameters: matrix size = 96×96×20; repetition time (TR)/echo time (TE) = 3000/15msec; voxel size = 0.312×0.312×1.2mm; slice thickness = 1mm; volume = 200. T2-weighted high-resolution anatomical scans were conducted using RARE pulse sequence with imaging parameters as follows: matrix size = 256×256×20; TR/TE = 4369/12msec; voxel size = 0.117×0.117×1mm; slice thickness = 1mm.

#### Pre-Scan Elevated Plus Maze

Prior to both PD28 and PD48 imaging, subjects were evaluated for anxiety-like behavior in the EPM to determine whether it could be predictive of rsFC alterations, and whether juvenile behavior/rsFC might be predictive of later adolescent outcomes in the same rat. Methods identical to Study 1.

#### Analysis of rsFC and Behavior

rsFC was measured using seed-based voxel-wise analysis, with BLA as the seeded region. rsFC between BLA and IL and PL regions of PFC were the focus of the present study. Data analyses were conducted using Analysis of Functional NeuroImages (AFNI_17.1.12; NIH), FMRIB software library (FSL, v5.0.9), and Advanced Normalization Tools (http://stnava.github.io/ANTs/). Resting-state Blood Oxygen-Level Dependent (BOLD) data were used for brain-tissue data extraction using 3Dslicer (https://www.slicer.org). Skull-stripped data were despiked to remove large signal fluctuations due to scanner and physiological artifacts. Slice-timing correction was conducted to correct data from interleaved slice order acquisition. Head motion correction was carried out using six parameters, with first volume as reference slice. Each subject was registered to a standard MRI Rat Brain Template (Ekam Solutions LLC, Boston, MA) using non-linear registration. In order to further reduce motion effects and physiological fluctuations, regressors comprised of six motion parameters, the average BOLD signal in white matter and ventricular regions, as well as motion outliers among all data volumes were fed into a nuisance regression model. Band-pass temporal filtering (0.01Hz-0.1Hz) and spatial smoothing (FWHM = 0.6mm) were performed on the residual data followed by signal detrending.

Whole brain voxel-wise Pearson’s correlation coefficients were calculated for each subject and were transformed for normality using Fisher’s z. Because this experiment was conceived to follow up on observed sex-specific effects on innervation measures, a priori hypotheses drove the use of a two-way mixed ANOVA within each sex (age as a within-subjects factor and stress as a between-subjects factor) in lieu of a three-way ANOVA. Therefore, two-way ANOVAs were conducted for each sex using AFNI 3dLME, with a false discovery rate curve computed. Post-hoc contrast effects focusing rearing condition (CON vs. ELS) at two levels of age (PD28 or PD48), as well as contrasts effects of age at two levels of rearing condition, were conducted where appropriate. Findings with a voxel-wise uncorrected p < 0.005, with a minimum cluster size of 30 voxels, was regarded as statistically significant.

Linear regressions were conducted to determine the relationship between EPM performance and correlation coefficients of BLA-seeded rsFC data for PL and IL in male and female rats. Since rats were tested at two time points, regressions were also performed to assess whether juvenile outcomes (behavior and/or rsFC) were predictive of later adolescent outcomes. Fisher’s r-to-z transformations were performed to determine any significant impact of sex or rearing on the strength of the relationships. All statistical analyses for behavior, as well as regression analyses to determine relationships between rsFC correlation coefficients and behavior, were performed in either SPSS v25 (IBM) or PRISM 8 (Graphpad).

## Results

### Study 1: Neuroanatomy and Axonal Innervation

#### BLA volume

A subset of tissue from each group was analyzed for BLA volume with Cresyl Violet. Average BLA volume for each age (PD28, PD38, PD48) was calculated using the Cavalieri probe in StereoInvestigator (n=16 per age). Two-way ANOVA for each age revealed no effects of sex or rearing on BLA volume (Supplemental Fig1). Thus, results were pooled across sex and rearing to determine BLA volume across development. One-way ANOVA showed a main effect of age (*F*_2,48_ = 31.74, *p* < 0.0001). Comparisons revealed increased BLA volume between PD28 and PD38 (*p* = 0.002), PD28 and PD48 (*p* < 0.0001); and PD38 and PD48 (*p* = 0.0002) (Fig2A).

#### BLA Bolus

Standard deviation of BLA volumes within each age deviated ~1% from mean volume, therefore percent BDA-filled BLA was calculated by dividing bolus volume in BLA in each animal by average BLA volume per age. Percent bolus outside BLA was calculated for each animal by dividing total bolus outside of BLA by total bolus volume.

Since age differences were detected in BLA volume, we examined whether amount of BLA filled differed with age. One-way ANOVA revealed no effect of percent BLA filled (*F*_2,88_ = 3.030, *p* = 0.053; small effect size: partial η2 = 0.06) (Fig2C), and no effect of percent bolus outside of BLA across age (*F*_2,88_ =0.133, *p* = 0.876) (Fig2D,E).

To ensure that percentage of bolus within BLA did not drive individual differences in PFC axonal innervation, linear regressions were used to determine correlations between percent of filled BLA volume and total innervation in the PFC, as well as the percent of BLA filled with tracer and total innervation in the PFC. These revealed no relationship between percent BLA filled and total innervation at any age (PD28: *R*^2^(28) = 0.064, *p* = 0.178; PD38: *R*^2^(27) = 0.004, *p* = 0.738; PD48: *R*^2^(30) = 0.057, *p* = 0.189). Additionally, linear regression analyses were used to investigate possible relationships between the total bolus volume (both within and outside of the BLA) and total PFC innervation, which revealed no significant correlation at any age (PD28: *R*^2^(28) = 0.062, *p* = 0.185; PD38: *R*^2^(27) = 0.006, *p* = 0.696; PD48: *R*^2^(30) = 0.090, *p* = 0.096) (Supplemental Fig2).

#### BLA-Derived Axonal Innervation

There were no differences in probe volume or mounted thickness (*p*> 0.1) between groups based on age, sex, or rearing; thus, all analyses are presented as collected without corrections (Supplemental Fig3).

#### PL innervation

A three-way ANOVA to determine effects of rearing, sex, and age on BLA-PL innervation revealed a main effect of rearing (*F*_1,79_ = 14.294, *p* < 0.0001), and a three-way interaction (*F*^2,79^ = 5.244, *p* = 0.007). Two-way ANOVA of male PL innervation showed a trending age x rearing interaction (*F*_2,42_ = 2.748, *p* = 0.076) with a moderate effect size (partial aηn2E=LS0-.1d1ri6v)e;nfoinllcorwe-auspe pinositn-nheorcvasthioonweadt PD38 compared to CON (*p* = 0.031). Increased innervation at PD38 compared to PD28 was observed in ELS (*p* = 0.044). Two-way ANOVA of female PL innervation showed a main effect of rearing (*F*_1,38_ = 14.21, *p* < 0.001), with post-hoc indicating more PL innervation in ELS compared to CON at PD28 (*p* = 0.009) and PD48 (*p* = 0.010). Graphs detailing comparisons, as well as representative photomicrographs of PL, can be seen in in Fig3A.

**Figure 3.**
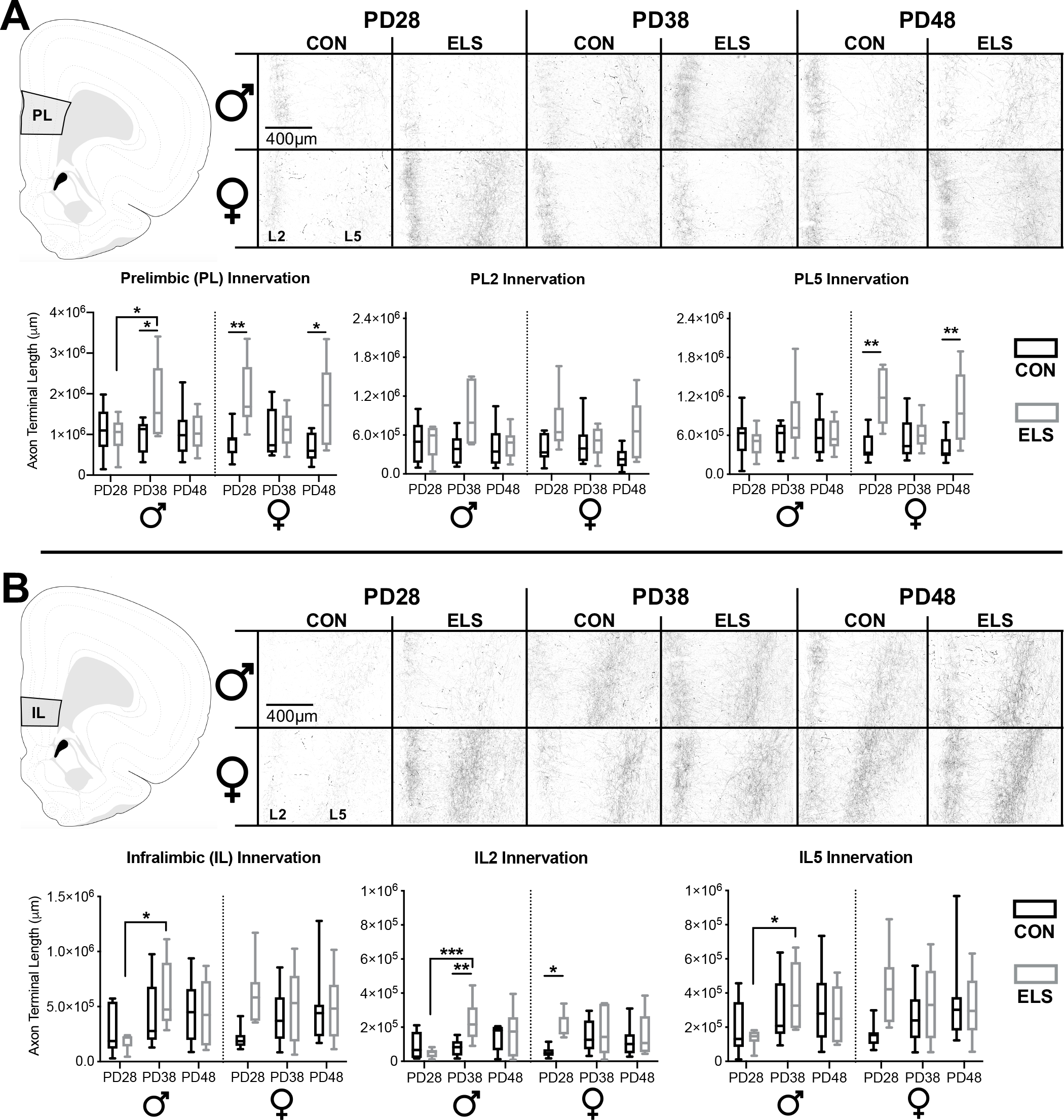
ELS leads to precocial BLA-PFC axonal innervation earlier in female than in male rats. BLA-PFC axonal innervation of anterogradely biotinylated dextran amine (BDA)-labeled fibers were visualized with diaminobenzidine (DAB) staining, quantified via unbiased stereology, and are displayed here as an estimate of total axon terminal length (μm) in male and female rats in both PL (**A**) and IL (**B**). Top panel (**A**) indicates PL quantification location, as well as the representative photomicrographs, with dark axonal fibers clearly observed in layer 2 (L2; left side of each photomicrograph) and layer 5 (L5; right side of each photomicrograph). In the PL there was consistent innervation across CON groups at all ages, with transient ELS-induced spikes of innervation occurring at PD28 and PD48 in female rats, and at PD38 in male rats. This pattern of innervation appears to be driven by PL5 axon terminal length, particularly in ELS females. Bottom panel (**B**) indicates IL quantification location, as well as representative photomicrographs similar to those seen in PL. In the IL, female rats showed a precocial pattern of axonal innervation by PD28 that was comparable to adolescents and young adults, while male rats didn’t show an ELS-driven increase in IL innervation until PD38, with these findings appearing to be driven by IL2 axon terminal length, particularly in ELS animals. Data is presented as a function of sex (male, female), age at brain collection (PD28, PD38, PD48), and rearing condition (CON (black bars), ELS (grey bars)), with lines inside each group bar indicating group mean, and lines outside of the bars indicating the maximum and minimum observed data points within that group. n = 6-9 per group. The left graph for both PL (**A**) and IL (**B**) displays collapsed L2 and L5 data for the entire quantified region, with the middle and right graphs displaying layer-specific data (alpha adjusted to 0.025 significance threshold to account for multiple comparisons in subsequent two-way ANOVAs). Photomicrographs were imaged at 10x mag. Atlas modified from Swanson (2018). \**p* < 0.05 (for IL/PL Innervation graphs); \**p* < 0.025 (for individual L2 and L5 innervation graphs); \*\**p* < 0.01; \*\*\**p* < 0.001

Three-way ANOVAs to delineate layer-specific effects in PL revealed a main effect of rearing in PL2 (*F*_1,79_ = 12.511, *p* = 0.001) and PL5 (*F*_1,79_ = 12.925, *p* = 0.001). There were significant rearing x sex x age interactions in PL2 (*F*_2,79_ = 4.655, *p* = 0.012) and PL5 (*F*_2,79_ = 4.699, *p* = 0.012). In PL5, a rearing x sex interaction was observed (*F*_1,79_ = 7.775, *p* = 0.007). Two-way ANOVAs were conducted for each sex. In PL2, males failed to show age (*p* = 0.202) or rearing (*p* = 0.043) effects that met the Bonferroni-corrected ɑ of 0.025. However, two-way ANOVA in females revealed a main effect of rearing (*F*_1,46_ = 5.344, *p* = 0.025) in PL2. Within PL5, males showed no significant effects or interactions (*p* > 0.10). However, two-way ANOVAs of PL5 innervation in females revealed a main effect of rearing (*F*_1,38_ = 17.760, *p* = 0.0001). Post-hoc revealed an effect of rearing with female ELS showing more PL5 innervation than CON at P28 (*p* = 0.002) and P48 (*p* = 0.007).

#### IL Innervation

Three-way ANOVA revealed no overall main effects or interactions in the IL as a whole, however a trending main effect of rearing was noted (*p* ≥ 0.07). However, since our a priori hypothesis was that the effects of ELS would be sex specific, we also performed separate 2-way ANOVAs for males and females. Two-way ANOVA in males showed a main effect of age (*F*_2,40_ = 4.355, *p* = 0.019), with increased IL innervation from PD28 to PD38 (*p* = 0.015) in ELS. Two-way ANOVA in females showed no main effect of age nor an age x rearing interaction. Graphs detailing comparisons, as well as representative photomicrographs of IL, can be seen in in Fig3B.

Three-way ANOVAs were conducted for IL2 and IL5. Adjusted ɑ was set to 0.025 to correct for multiple comparisons. In IL2, a main effect of rearing was evident (*F*_1,79_ = 5.938, *p* = 0.017), with no main effects or interactions in IL5 (*p* > 0.10). Two-way ANOVAs were conducted for each sex and sub-region. Males displayed a main effect of age (*F*_2,39_ = 4.411, *p* = 0.019) in IL2, and a rearing x age interaction (*F*_2,39_ = 4.521, *p* = 0.017). Male ELS showed more IL2 innervation compared to CON at PD38 (*p* = 0.007). Developmentally, male ELS had more innervation at PD38 than PD28 (*p* < 0.001). In females, there was a trending main effect of rearing (*F*_1,38_ = 4.791, *p* = 0.035) with a moderate effect size (partial η2 = 0.126), with more IL2 innervation in ELS than CON at PD28 (*p* = 0.037). Two-way ANOVA on male IL5 showed a trending main effect of age (*F*_2,40_ = 3.541, *p* = 0.038; partial η2 = 0.178), with post-hoc revealing more innervation at PD38 than PD28 in ELS (*p* = 0.048). Two-way ANOVA on female IL5 showed no main effects (*p* > 0.1) or interaction (*p* = 0.096).

#### EPM

Three-way ANOVA comparing time spent in the open arms showed an interaction of age x rearing (*F*_2,224_ = 3.06; *p* = 0.049) and of sex x rearing (*F*_2,224_ = 5.98; *p* = 0.015) (Fig4A). Two-way ANOVAs revealed a main effect in females of rearing (*F*_1,128_ = 12.69, *p* = 0.0005), with ELS females spending less time in the open arms than CON at PD38 (*p* = 0.0176). No effects of treatment or age were observed in males (*p* > 0.40). Arm crossings and head dips were also analyzed. No interactions or main effects of age, sex, or rearing were found with a 3-way ANOVA on head dips (Fig4B), however an age x sex interaction (*F*_2,242_ = 6.73; *p* = 0.001) and follow-up 2-way ANOVA revealed a main effect of age in males (main effect of age: *F*_2,114_ = 7.22; *p* = 0.007), with fewer arm crosses at PD48 than at PD28 (*p* = 0.0006) or PD38 (*p* = 0.0203) (Fig4C).

**Figure 4.**
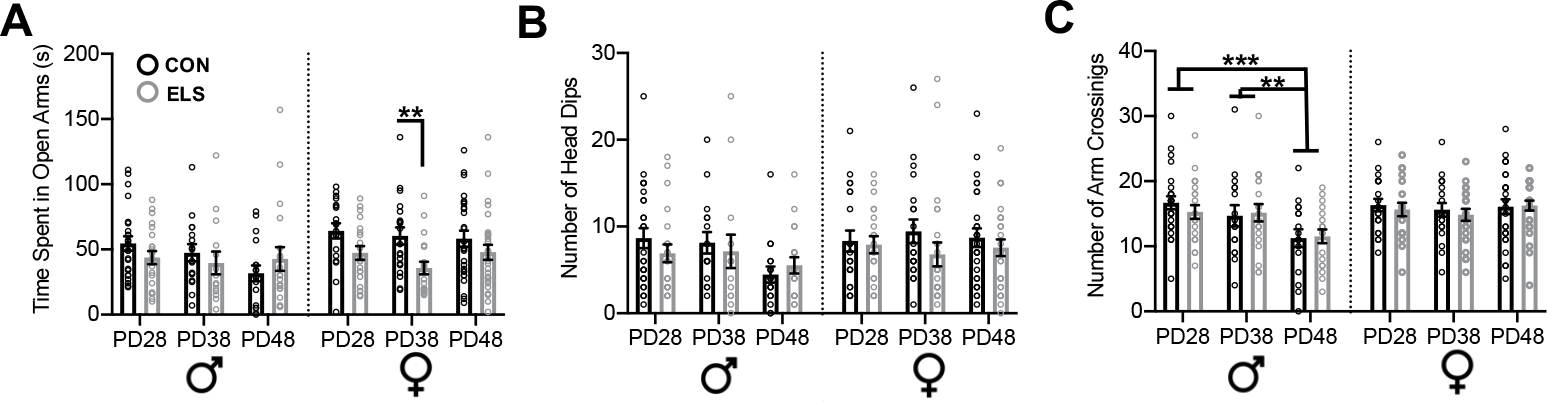
Anxiety-like behavior following ELS is sex-and age-specific. Anxiety-like behavior was assessed in the elevated plus maze (EPM) in both male and female CON (black bars) and ELS (grey bars) groups. Anxiety-like behavior in the EPM was assessed via time spent in open arms (seconds; **A**), number of head dips (**B**), and number of arm crossings (**C**). Female, but not male, rats showed more anxiety-like behavior (measured via less time spent in open arms), exclusively at PD38 if they had been exposed to ELS (**A**). Analysis of the number of arm crossings (**C**) revealed that only males showed a significant effect of age, with PD48 males showing less arm crossings (indicative of increased anxiety) than those at PD28 and PD38. Each circle indicates the data from a single animal, and data shown is for behavior collected from all subjects, including those that did not meet neuroanatomical inclusion criteria. *n* = 16-28 per group. \*\**p* < 0.025; \*\*\**p* < 0.001

#### Relationships Between Innervation and Behavior

Results from all analysis of correlation between behavior on the EPM and connectivity measures are shown in Table 1. Here we discuss significant correlations, which are illustrated in Fig5A-D. Fisher’s r-to-z transformations revealed an impact of sex, but not rearing, on the strength of some relationships between innervation and behavior (Table 1). At PD28, higher IL innervation in females was correlated with less time spent in the open arms (*R*^2^(14) = 0.478; *p* = 0.009) (Fig5B). At PD38, males showed a similar relationship between less time in the open arms and higher PL innervation (*R*^2^(13) = 0.266; *p* = 0.049) (Fig5C) and higher IL innervation (*R*^2^(13) = 0.260; *p* = 0.05) (Fig5D). No relationships were seen at PD48 (Table 1).

**Figure 5.**
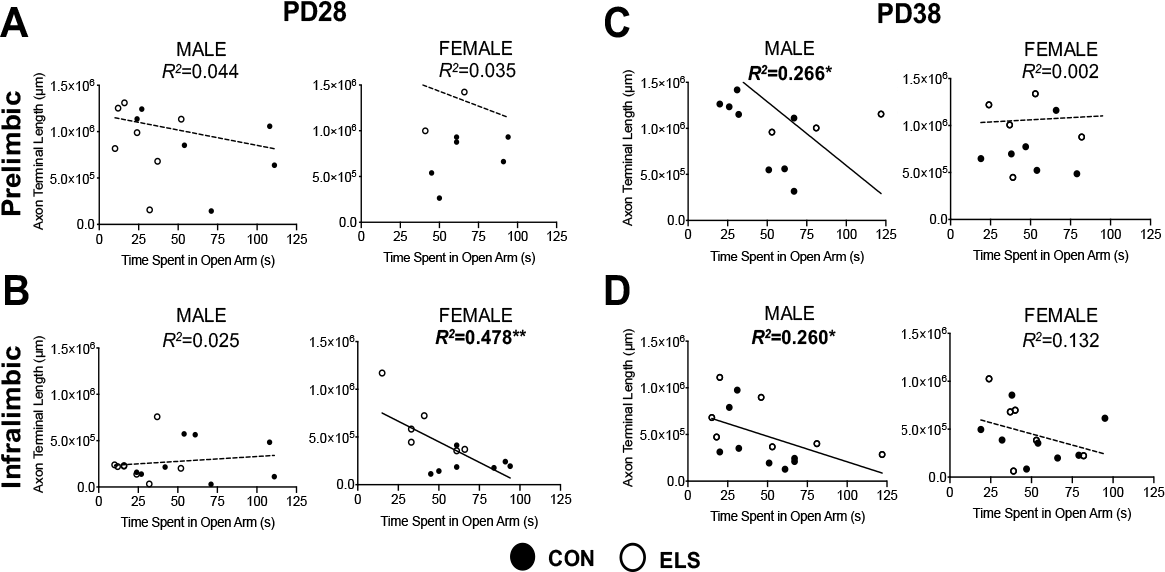
More BLA-derived axonal innervation into the PFC is correlated with increased anxiety-like behavior in a sex- and age-dependent manner. Results of linear regression analyses to determine the relationship between BLA-PFC innervation (measured as an estimate of BDA-labeled axon terminal length) and performance in the EPM (time spent in open arms (seconds) as an index of anxiety-like behavior). At PD28, there were no significant correlations between axon terminal length in the PL and time spent in open arms of the EPM (**A**) in either males or females, regardless of rearing condition. However, females, but not males, showed a significant correlation (**B**) suggesting that more axonal innervation of the IL was related to increased anxiety-like behavior (via decreased time spent in open arms). Conversely, at PD38, females no longer show behavior correlated with innervation in either PL (**C**) or IL (**D**), whereas males generally show that increased axonal innervation is correlated with increased anxiety in both of these regions (**C**; **D**). Regression lines are reflective of the overall relationship across all groups (CON and ELS), and individual subjects from anatomically included cases are signified by either solid black circles (CON) or open circles (ELS). Bold lines/regression statements indicate a significant correlation. *n* = 6-9 per group. \**p* < 0.05; \*\**p* < 0.01

**Table 1.**
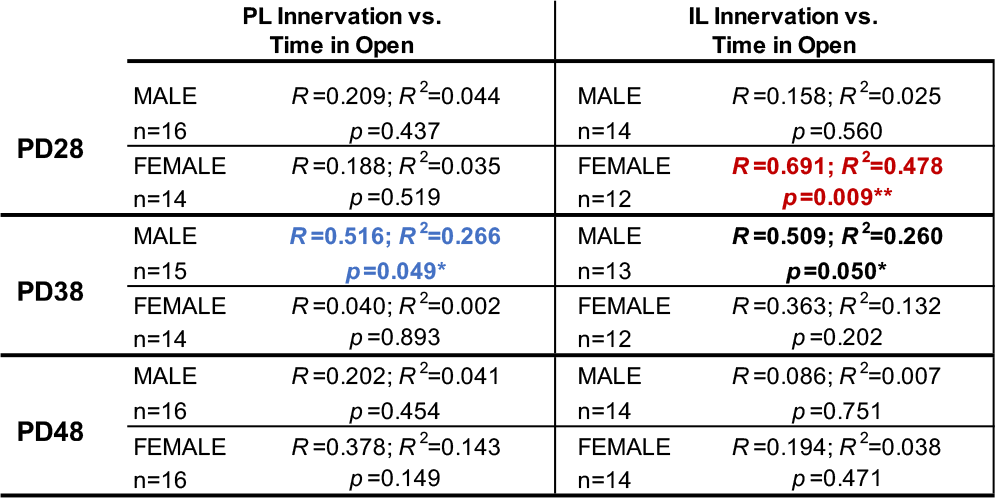
Relationships between BLA-PFC innervation and anxiety-like behavior across development. Within the PL, there was a significant correlation between BLA innervation and time spent in open arms (highlighted in bold) of the EPM exclusively in males at PD38. Fisher’s r-to-z transformation also revealed a significant effect of sex (signified by blue bold typeface). There was also a significant correlation between BLA innervation of the IL and time spent in open arms of the EPM in males at PD38 (bold type). Within the IL, female showed a significant correlation between BLA-PFC innervation and time spent in open arms exclusively at PD28, and Fisher’s r-to-z transformation also revealed a trending effect of sex (signified by bold red typeface) at this age. Since there was no significant effect of rearing observed, data was collapsed across rearing groups for correlational analyses presented here. n’s = 12-14/age group. **Bold** (*p* < 0.05); significant correlation without meeting criterion for significant sex effect Bold and Blue (*p* < 0.05); significant correlation and significant sex difference Bold and Red (*p* = 0.06); significant correlation and trend-level sex difference

### Study 2 Results: rsFC

#### Males

Two-way ANOVA evaluating BLA-PL rsFC (analyses of these regions were seeded to the BLA; Fig6A-B) with age and rearing revealed no main effect of ELS, but a significant main effect of age (*F* = 49.9) and two regions with an age x rearing interaction in PL that met threshold criteria (*F* = 15.8; *F* = 28.6). Post-hoc showed reduced strength of BLA-PL rsFC from PD28 to PD48 in both CON (Z = −3.82) and ELS (Z = −3.26). There was no difference in rsFC between CON and ELS at PD48; however, at PD28 CON showed stronger BLA-PL connectivity than ELS (Z = −4.13; Fig6C). A similar 2-way ANOVA evaluating BLA-IL rsFC revealed a significant main effect of age (*F* = 49.9), with no post-hoc differences.

**Figure 6.**
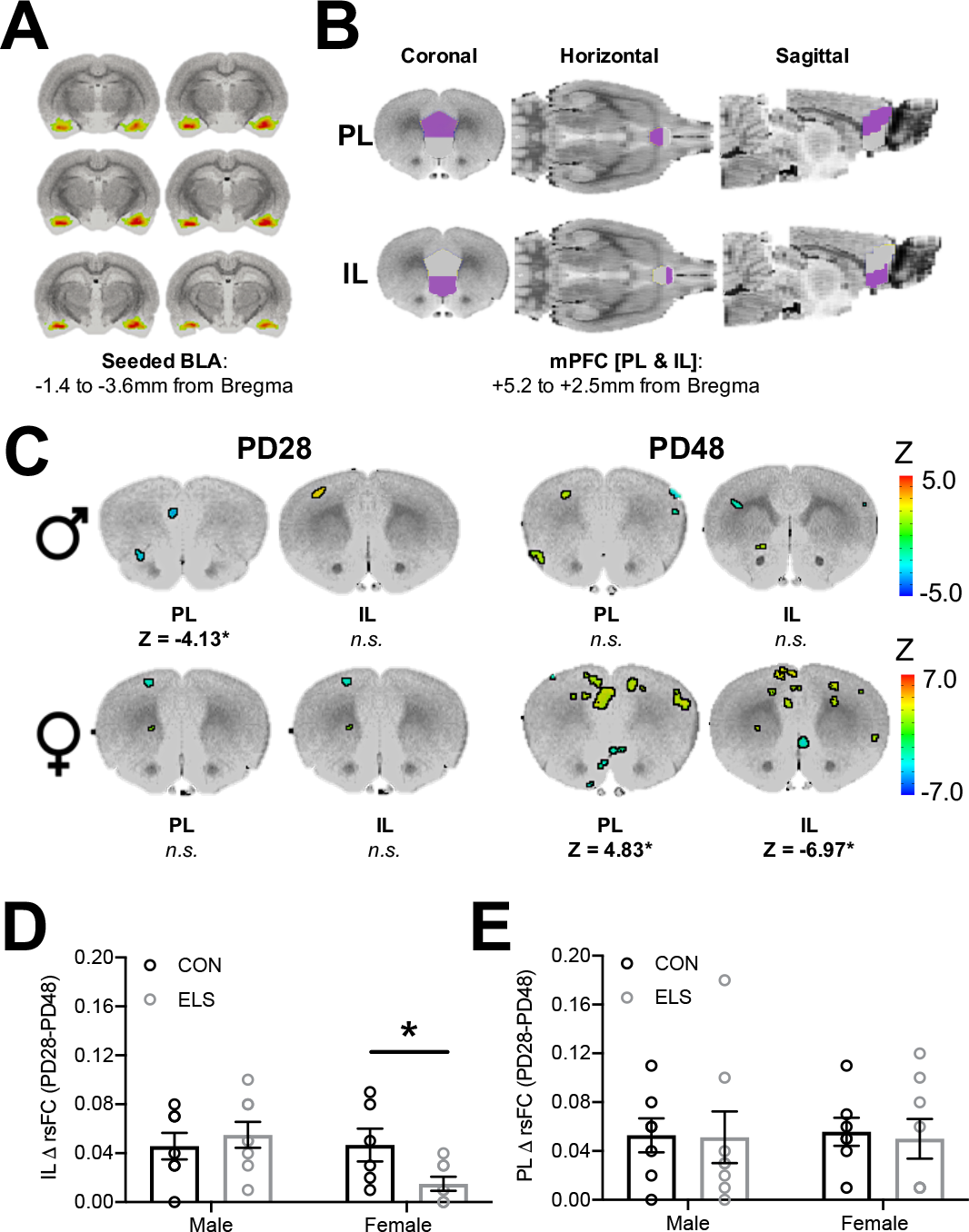
Effects of ELS on BLA-PFC rsFC are sex-specific and endure in females. Resting state functional connectivity (rsFC) was assessed in a within-subjects manner across development in male and female rats with a history of either CON or ELS rearing with the basolateral amygdala (BLA) as the seeded region (**A**). The rsFC between the BLA and the mPFC – specifically the prelimbic (PL) and infralimbic (IL) cortices (**B**) – was assessed. (**C**) shows results from comparative analyses comparing CON and ELS groups at each age point (PD28 or PD48) in both male and female groups. Colored regions correspond to computed Z values and indicate specific regions within either the PL or IL that met criteria for significance with a minimum cluster size of 30 voxels. Generally, male rats show no group effects of ELS, with the exception of a finding of decreased BLA-PL rsFC in ELS compared to CON groups at PD28. Female rats showed no effect of ELS at PD28, but show striking differences in rsFC in both the PL and IL at PD48 (**C**). Female, but not male, rats exposed to ELS showed a lack of typical maturation of the IL, evidenced by significantly reduced ΔrsFC from PD28 to PD48 compared to CON females (**D**). Conversely, there were no group changes in ΔrsFC observed within the PL (**E**). n = 7-8 per group. *p < 0.05

#### Females

Two-way ANOVA (age x rearing) for BLA-PL revealed a main effect of age (*F* = 13.76) and rearing (*F* = 16.10), and an interaction (*F* = 12.65). Overall, BLA-PL rsFC decreased from PD28 to PD48 (Z = −3.6), while ELS females displayed stronger connectivity compared to CON (Z = 3.5). Post-hoc showed no difference between CON and ELS at PD28, though a significant difference at PD48 when ELS had stronger BLA-PL connectivity than CON (Z = 4.83; Fig6C). CON connectivity decreased from PD28 to PD48 (Z = −3.09). Conversely, ELS BLA-PL connectivity increased from PD28 to PD48 (Z = 3.46).

Two-way ANOVA for BLA-IL rsFC showed no main effect of age or rearing, though an interaction was observed (*F* = 28.74). Overall, BLA-IL rsFC increased from PD28 to PD48 (Z = 3.35), and CON generally had weaker connectivity compared to ELS (Z = −3.79). Post-hoc revealed that, at PD48, ELS showed less BLA-IL connectivity than CON (Z = −6.97; Fig6C), and CON showed increased BLA-IL connectivity from PD28 to PD48 (Z = 4.47), with no significant change seen in ELS.

After noting what appeared to be a lack of typical BLA-PFC rsFC maturation in ELS-exposed females, we then analyzed the effects of sex and rearing on the magnitude of change in BLA-IL rsFC and BLA-PL rsFC between PD28 and PD48. 2-way ANOVA of BLA-IL rsFC revealed a moderate effect size for a trend-level sex x rearing interaction (partial η2 = 0.167; *F*_1,25_ = 4.09; *p* = 0.054), and post-hoc revealed that females exposed to ELS displayed less PD28-PD48 change compared to CON females (*p* = 0.033) (Fig6D). In contrast, no effect of rearing was observed for the magnitude of change in BLA-PL rsFC (*p*=0.78) (Fig6E).

#### Relationships Between Innervation and Behavior

Results from all regression analyses of the relationship between behavior on the EPM and functional connectivity are shown in Tables 2 and 3. Fisher’s r-to-z transformations revealed an impact of both sex and rearing on relationships between rsFC and behavior at PD28. Significant correlations revealed that in PD28 females exposed to ELS, lower BLA-IL rsFC predicted less time spent in the open arms (*R*^2^(8) = 0.852; *p* = 0.001) (Fig7A); notably, this relationship is juxtaposed with our finding that higher axonal innervation at the same age predicted less time spent in the open arms in all females (Fig5B). Our longitudinal design further allowed the analysis of early rsFC with later adolescent behavior. In all females, lower BLA-IL rsFC at PD28 also predicted less time spent in the open arms twenty days later at PD48 (*R*^2^(7) = 0.405; *p* = 0.011), suggesting an enduring and predictive relationship in females (Fig7B). This enduring relationship in females was also seen in a significant relationship between lower BLA-IL rsFC at PD48 and less time in the open arms at the same age (Fig 7C; *R*^2^(13) = 0.303; *p* = 0.041).

**Figure 7.**
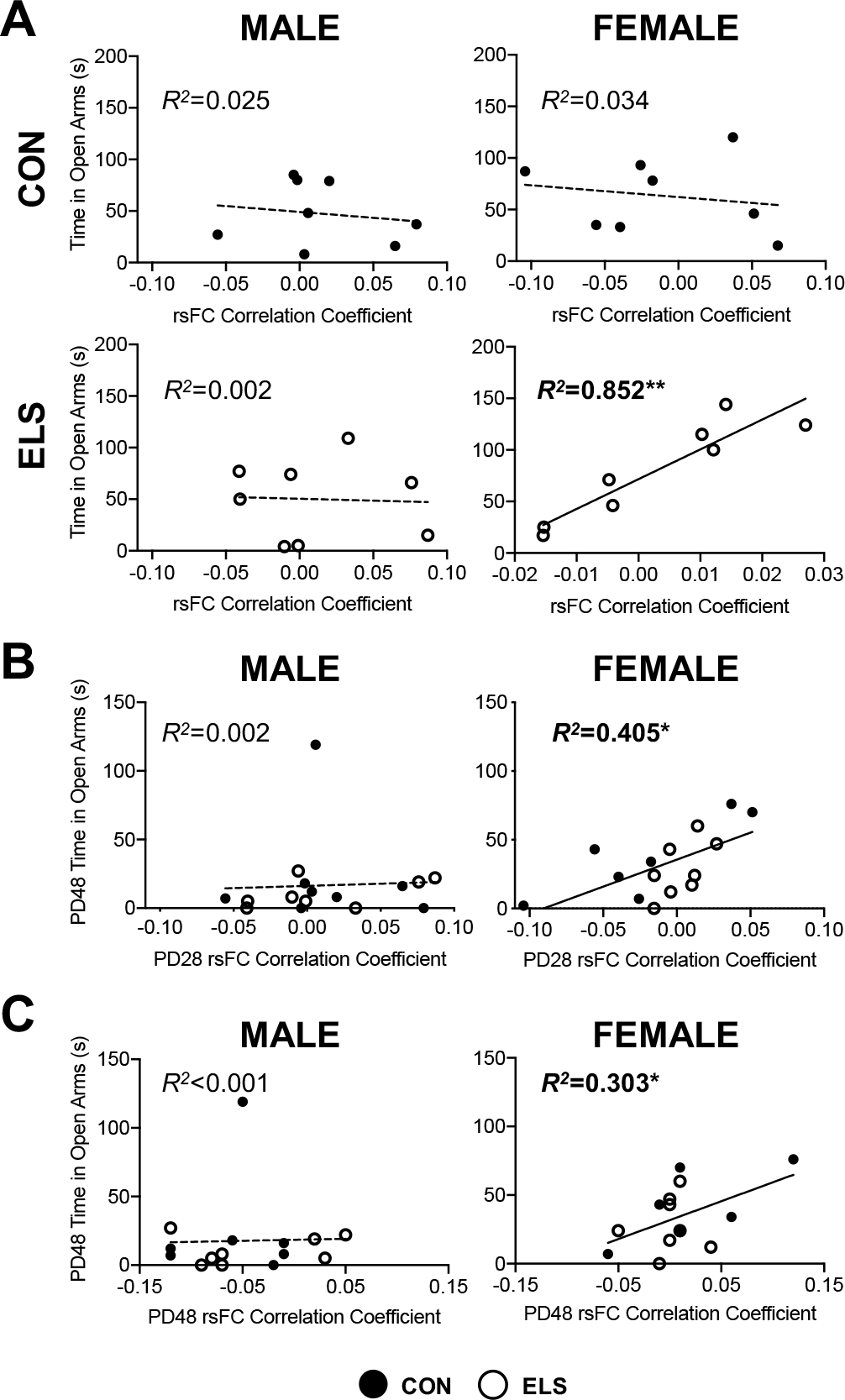
ELS significantly impacts the relationship between BLA-IL rsFC and anxiety-like behavior only in females; with PD28 rsFC predictive of PD48 behavior. Results of linear regression analyses to determine the relationship between BLA-IL rsFC correlation coefficients and performance in the EPM (time spent in open arms (seconds) as an index of anxiety-like behavior) are shown for PD28 in (**A**). Only females with a history of ELS showed a significant correlation, with higher rsFC correlation coefficients correlating with more time spent in the open arms of the EPM. As this data was conducted over development in a within-subjects manner, we could further explore predictive relationships between these variables. In line with other rsFC data described here, only female rats showed an overarching predictive effect of early/juvenile (PD28) rsFC correlation coefficients on later (PD48) behavior (**B**). Indeed, females with a higher rsFC correlation coefficients at PD28 exhibited less anxiety-like behavior (as evidenced by increased time spent in open arms). Individual data points for each animal can be seen on the graphs, with black circles representing CON cases, and open circles representing ELS cases, with solid regression lines indicative of significant correlations. *n* = 7-8 per group. \**p* < 0.05; \*\**p* < 0.01

**Table 2.**
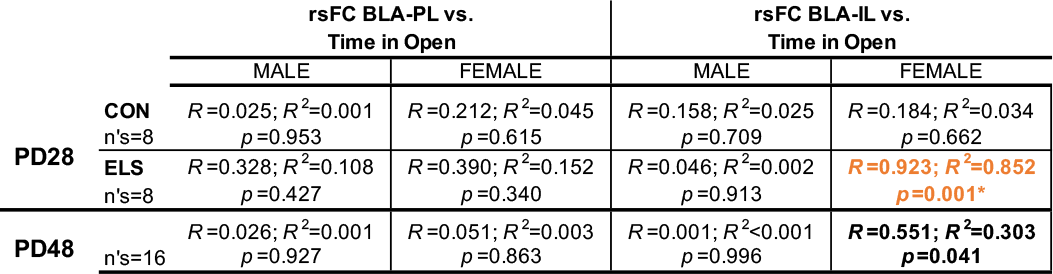
Relationships between BLA-PFC rsFC and anxiety-like behavior. Significant effects of rearing were observed at PD28, but not PD48, therefore analyses were performed separately in CON and ELS at PD28. Significant correlations were only observed between IL and time spent in the open arms of the EPM, and only in females. At PD28, ELS females showed a significant correlation between BLA-IL rsFC and time spent in open arms, and this was also characterized by a significant effect of sex determined by Fisher’s r-to-z transformation. At PD48, only females (across both rearing groups) show a significant correlation between BLA-IL rsFC and time spent in open arms. n’s = 8 for PD28 groups; n’s = 16 for PD48 groups. Bold (p < 0.05); significant correlation without meeting criterion for significant sex effect Bold and Orange (p < 0.05); significant correlation and significant effect of sex Asterisk (*) designates a significant effect of rearing (p < 0.05) within each sex.

**Table 3.** Predictive relationships between PD28 BLA-PFC rsFC and PD48 anxiety-like behavior. Within the IL, there was a significant correlation between female PD28 rsFC correlation coefficient and later PD48 time spent in open arms in the EPM, thereby suggesting a predictive value of early rsFC measures and later behavior. Since there were no effects of rearing observed, data was collapsed across rearing groups. Bold (*p* < 0.05); significant correlation without meeting criterion for significant sex effect

| PD48 Time Spent in Open vs. |  |  |  |
| --- | --- | --- | --- |
| PD28 rsFC BLA-PL |  | PD28 rsFC BLA-IL |  |
| MALE | $R=0.354$; $R^2=0.125$<br>$n=16$<br>$p=0.178$ | MALE | $R=0.049$; $R^2=0.002$<br>$n=16$<br>$p=0.856$ |
| FEMALE | $R=0.428$; $R^2=0.175$<br>$n=15$<br>$p=0.107$ | FEMALE | **$R=0.637$; $R^2=0.405$**<br>**$n=15$**<br>**$p=0.011^*$** |

## Discussion

This work describes sex-specific neuroanatomical etiology and corresponding functional maturation of the BLA-PFC circuit following ELS. The data reveal newly uncovered aberrations to PFC innervation that can be interpreted in the context of atypical behavior and FC that has been observed in humans (Gee et al., 2013a; Philip et al., 2013; Teicher et al., 2016; Thomason et al., 2015) and in animals (Johnson et al., 2018; Yan et al., 2017). We observed sex- and age-dependent effects on BLA innervation in PL and IL regions of the PFC following ELS in a rat model of caregiver deprivation. Our findings suggest that females may be particularly vulnerable to neuroanatomical consequences of early adversity, with innervation effects seen earlier in females compared to a later effect observed in males. This is in line with human studies describing sex-dependent effects of early adversity where female participants appear to exhibit more severe adolescent and later-life consequences, especially with regard to affective disorders (Humphreys et al., 2015).

Our data indicate that BLA-IL innervation increases through adolescence in a bilaminar manner across development, with the most dramatic increases following ELS in both males and females. Importantly, these main effects of age support previous work in male rats (Cunningham et al., 2002) and now characterize a similar trajectory in female rats. While IL innervation showed a developmental increase through adolescence, PL innervation appeared to reach adult-like levels in both sexes earlier than was captured presently (Bouwmeester et al., 2002; Van Eden and Uylings, 1985). Following ELS, however, our data indicate that aberrant effects on innervation were apparent at PD28 in females, but not until PD38 in males; this finding was observed across both the PL and IL. However, while ELS conferred a similar age-dependent influx of BLA axonal innervation in both regions, the long-term neuroanatomical alterations were different depending on region examined. Indeed, in PL we saw a transient effect of ELS that was age- and sex-dependent. In IL, however, there was evidence of precocial maturation, such that ELS induced BLA-IL innervation that was comparable to more mature (PD48) patterns in female juveniles (PD28) and male early adolescents (PD38). In fact, throughout the mPFC we observed that both male and female rats showed an unexpected transient spike in innervation that appeared to resolve at the following developmental timepoint. It is possible that reversal of the innervation seen at earlier time points may be due to pruning mechanisms (Koss et al., 2013) that typically occur in adolescence, and these mechanisms may serve to mediate excess innervation (Rakic et al., 1994; Riccomagno and Klodkin, 2016; Spear, 2000). Prior work has shown a peak in BLA-PL connectivity, specifically at PD30 (Pattwell et al., 2016), that may contribute to the present findings of transient ELS-exacerbated hyper-innervation. However, this does not explain why innervation to the IL spiked at PD28, was reduced to CON levels at PD38, and then spiked again by PD48 in female ELS. It is possible that the temporary (one week) isolation following surgery may have acted as a secondary stressor acting in conjunction with pubertal changes to produce a resurgence of this ELS-specific phenotype (Andersen and Teicher, 2008; Tzanoulinou and Sandi, 2016). Relatedly, surgery itself during peripubertal development compared to other time periods may have differentially interacted with ELS.

Importantly, the ages at which ELS-exposed females and males first displayed increased BLA-IL innervation (PD28 and PD38, respectively) were the ages at which higher innervation correlated with higher anxiety-like behavior (less time spent in open arms of the EPM). Since monosynaptic input from the BLA to the medial PFC drives aversion to anxiogenic stimuli as measured in the EPM (Felix-Ortiz, 2016), these data suggest that hypertrophic effects of ELS at the BLA-IL circuit lead to heightened anxiety-like behavior in ELS-exposed animals. However, we were surprised that group-wise comparisons only showed higher anxiety-like behaviors in ELS females at PD38, but not at PD28 when innervation was increased (Fig4). Group comparisons also failed to reveal higher anxiety-like behaviors in ELS-exposed males at any age. Our group and others have previously observed increased anxiety-like behaviors in male and female rats following ELS (e.g., Ganguly et al., 2015, Holland et al., 2013, Jin et al., 2018), therefore further work will reveal whether more sensitive assays can more reliably demonstrate anxiety-like behavior in ELS-exposed rats. Here we demonstrate that, regardless of mean differences, ELS experience can forge a relationship between heightened innervation and anxiety-like behavior at distinct developmental time-points that could suggest mechanistic ties between early hypertrophy of inputs and behavior.

Transient effects of ELS on innervation and behavior are notable because adolescent perturbations have been shown to have lasting consequences on later life function. Multiple lines of evidence from both clinical and animal studies suggest that physiological or experiential anomalies during critical periods of development can program the central nervous system for susceptibility or resilience to future environments (Andersen, 2003; Nederhof and Schmidt, 2012). One example is seen with an animal model using prenatal treatment with the mitotoxin MAM, which produces adolescent-specific corticolimbic and mesolimbic dysfunctions that drive a hyperresponsivity to stress with anxiety-like and psychosis-like behavior. If conversion to the dysfunctional phenotype in adolescence is prevented by relieving stress in adolescence, MAM-treated animals will still be more susceptible to affective dysfunction in adulthood (Gomes et al., 2019). Moreover, ELS reportedly leads to adolescent alterations in PFC NMDA receptor subunit composition that regulate anxiety-like behaviors in males (Ganguly et al., 2015); future work will reveal whether increased glutamatergic input from the BLA drives these receptor changes in the PFC. Since the PFC can serve reciprocally as a modulator of BLA activity (Rosenkranz and Grace, 2002), it is therefore possible that transient innervation changes lead to long-term receptor alterations and subsequent changes to the efficiency of BLA-PFC functional connectivity. Together, the early increases in innervation after ELS reported here may perturb normal critical period maturation, leading to late-onset effects on functional connectivity and affective dysregulation.

The results from the rsFC analyses in females supported the idea that early maturation of BLA-IL innervation disrupted the later functional relationship between these regions (Tottenham and Galván, 2016). Specifically, females exposed to ELS displayed dampened maturation of BLA-IL rsFC compared to female CON; while no effects of rearing were apparent at PD28, ELS female rats failed to display the increase in BLA-IL rsFC by PD48 that was observed in female CON. This is in line with previous work showing that BLA-mPFC connectivity is grossly unchanged at pre-weaning following a limited bedding model (Guadagno et al., 2018), while rsFC later in life can be altered following ELS (Nephew et al., 2017). Interestingly, increased [more mature] rsFC BLA-IL connectivity in female ELS at PD28 correlated with decreased anxiety-like behavior at that age as well as 20 days later. This suggests that more mature connectivity in female juveniles may confer enduring behavioral resilience, which is consistent with previous reports that dampened connectivity is associated with increased anxiety in adolescence (Kim et al., 2011; Nooner et al., 2013). While ELS exposure in males appeared to result in lower juvenile (PD28) rsFC between the BLA and the PL, no effects on the magnitude of change between PD28-PD48 were noted, therefore altered maturation per se was not observed. Interestingly, ELS affected male BLA-PL rsFC at an age that preceded any ELS-attributable increased innervation, suggesting that the reciprocal connectivity between the two regions was altered without the influence of aberrant amygdalofugal innervation.

ELS experience resulted in dampened maturation of BLA-IL rsFC in females, in contrast to the precocial maturation we observed in axonal innervation of the IL from the BLA, as well as the accelerated maturation of task-based FC reported following childhood maltreatment in humans (Gee et al., 2013a). While BLA-mPFC rsFC in humans has been observed to increase towards more positive connectivity through adolescent development (Gabard-Durnam et al., 2014), task-based FC during fearful face presentations declines from positive connectivity to a negative connectivity (Gee et al., 2013b). This task-based FC was further found to reach mature (more negative) levels in previously institutionalized children (Gee et al., 2013a). In contrast, girls who express higher basal cortisol levels at 4.5 years display lower rsFC between the ventromedial PFC and the amygdala at 18 years (Burghy et al., 2012), corroborating our current findings in PD48 females (Fig6). rsFC reflects an intrinsic alternate resonance between different brain areas connected at large scale, in contrast to the more isolated activation of the corresponding brain areas during a task (Rasero et al., 2018). Therefore, it appears that early hyperinnervation to the PFC may alter the stability of the functional BLA-PFC network as it develops, leading to increased anxiety when innervation arises and disrupted maturation of rsFC; how these effects in rats relate to task-based FC is currently unknown. It is also possible that early maturation of BLA-PFC projections may overwhelm the slow-maturing reciprocal projections to the BLA that dampen anxiety-related circuit activity in mature brains (Arruda-Carvalho et al., 2017; Selleck et al., 2018).

The present findings support previous work indicating the distinct functional relationship between the BLA and PL and IL regions of mPFC (Calhoon and Tye, 2015), with PL connectivity related to anxiogenic effects (Felix-Ortiz et al., 2016) and IL connectivity promoting anxiolytic effects (Maroun et al., 2012). Furthermore, these findings are in line with the idea that altered neuroanatomy and functionality of PL and IL, along with BLA, likely have reciprocal effects on one another, further contributing to the exacerbation of ELS-induced anxiety-like phenotypes in adolescence and early adulthood (Likhtik and Paz, 2015; Zimmermann et al., 2019).

Taken together, maternal separation in rats was found to disrupt normative development of both anatomical (axonal innervation) and functional (rsFC) connectivity within a circuit regulating emotional processing, with sex-specific effects and evidence of resilience in individuals with precocial maturation of rsFC. These findings have implications for intervention based on experience, age, and sex – three functionally interactive factors that uniquely define risk in every individual.

## Acknowledgements

The authors would like to sincerely thank Dr. Jens Foell (Florida State University) and Dr. Michael Rohan (McLean Hospital) for their feedback and suggestions for rsFC figures for this manuscript.

