## Supplemental Figures 1-3 for "Too much, too young? Altered corticolimbic axonal innervation and resting state functional connectivity suggests sex-dependent outcomes in a rat model of early life adversity"

### Supplemental Figures & Figure Legends

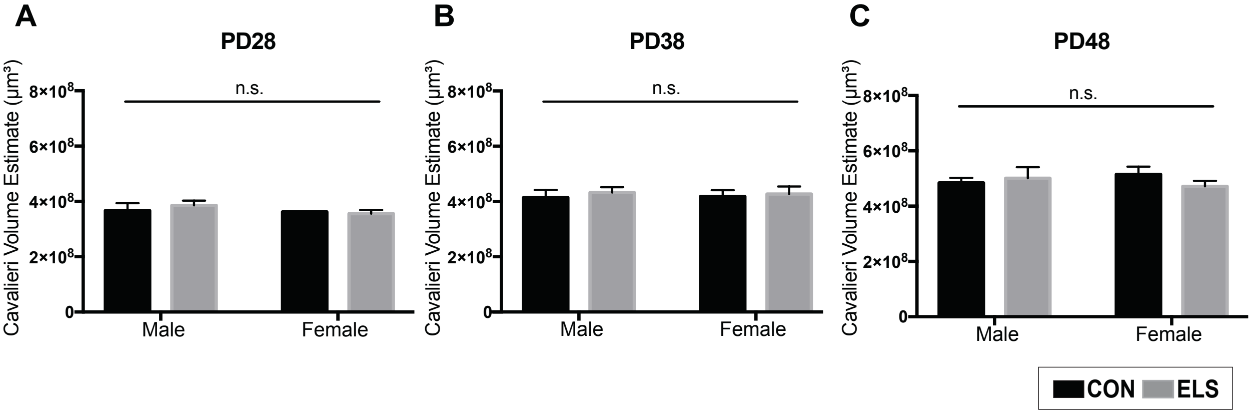


**Supplemental Figure 1: No effects of sex or rearing condition on BLA volume.** There were no significant effects of sex or rearing on basolateral amygdala (BLA) volume as estimated via Cavalieri probe across the rostral-caudal extent of the structure in PD28 (**A**), PD38 (**B**), or PD48 (**C**) rats. n=4 per group. As such, data presented in panels **A**, **B**, and **C** were collapsed across sex and rearing condition for each age group to determine average BLA volume (n=16 per age).
n.s. (non-significant)

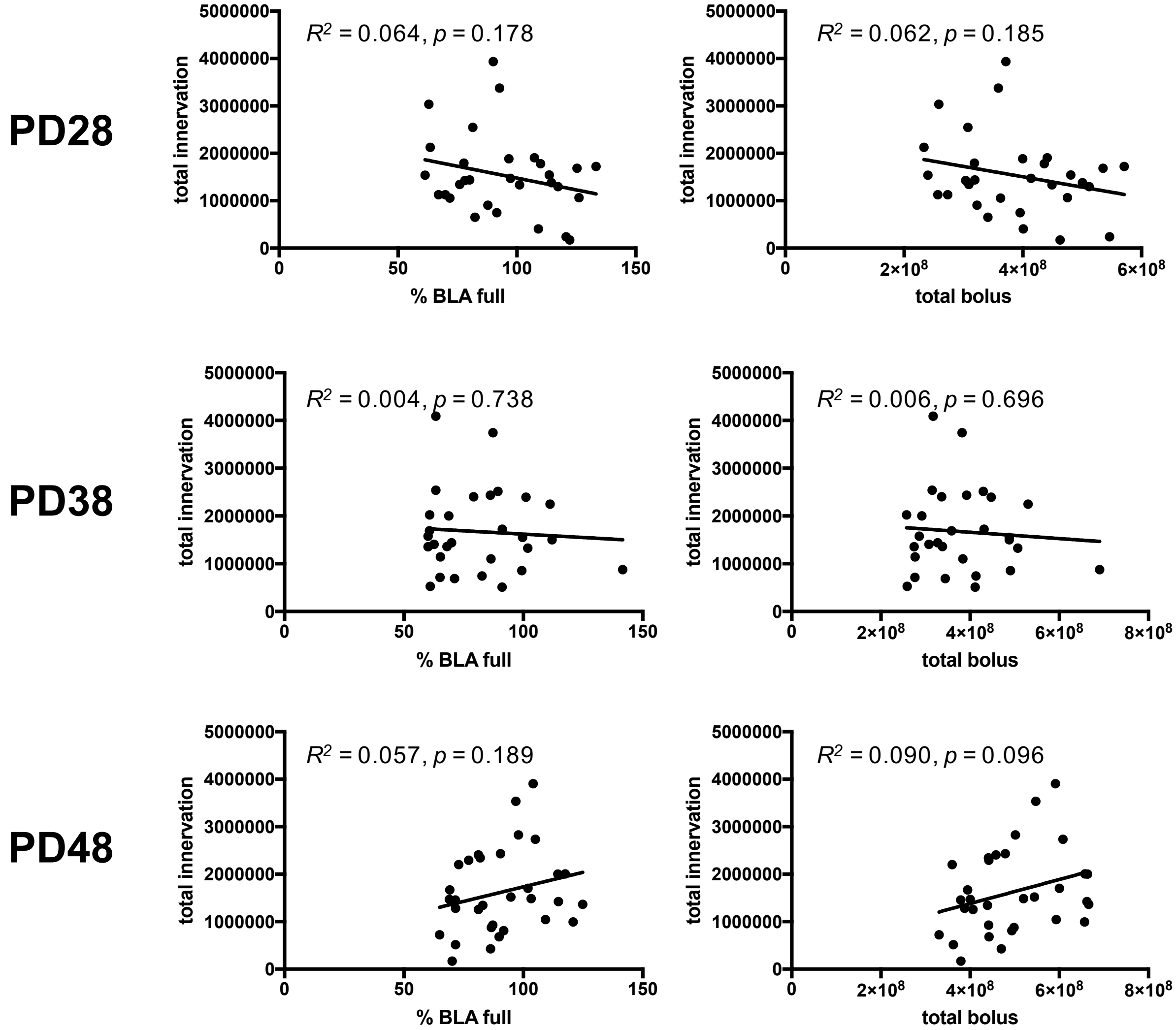


**Supplemental Figure 2: PFC innervation is not driven by percentage of BLA filled or bolus size in included cases.**
All included cases for each age (PD28, PD38, and PD48) were collapsed across sex and rearing condition to determine whether the percentage of BLA that was filled, and/or the total bolus size (both within and outside of the BLA structure) was related to total PFC axonal innervation in included cases (n=29-32 per age). There were no observed relationships between these measures at any age, suggesting no need for correcting innervation based on bolus characteristics measured. Bolus size and location (i.e. % in and outside of BLA structure) were determined via Cavalieri probe volume estimate across the rostral-caudal extent of the biotinylated dextran amine (BDA) anterograde tracer bolus.

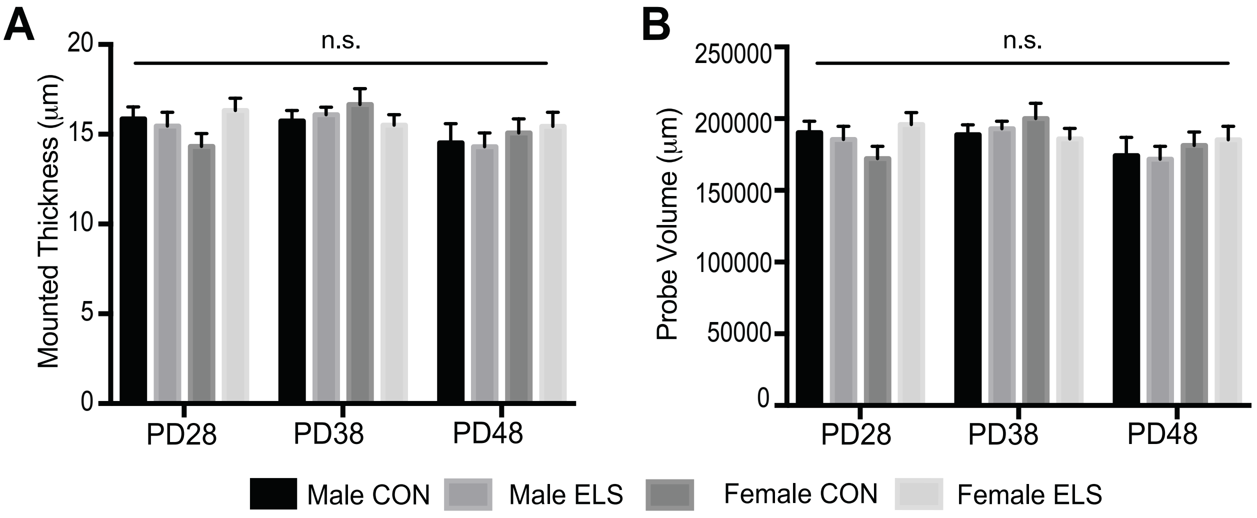


**Supplemental Figure 3: No differences in mounted thickness or probe volume.** Mounted thickness (⌠m) was assessed across the Z plane for each quantified tissue section for each animal, and this measure was averaged for each animal, and an average tissue thickness for each group within each age was computed (**A**). There were no significant differences in mounted thickness across groups at any age, as well as no significant differences as a function of age. The average total probe volume (PL+IL) for StereoInvestigator analysis of BLA-PFC axon fiber length for each brain region per animal was also computed for each age (**B**), and no significant differences were found. n=6-9 per group.
n.s. (non-significant)
